# Natural variation in the oxytocin receptor gene predicts social observation in female prairie voles

**DOI:** 10.1101/2025.11.03.686389

**Authors:** Shang Lin Tommy Lee, Xinyuan Max Cao, Sena Agezo, Arjen J. Boender, Caroline Bowen, Zachary V. Johnson, Larry J. Young, Gordon J. Berman, Robert C. Liu

## Abstract

Genetic variation in the oxytocin receptor gene (*Oxtr*) has been linked to differences in brain OXTR expression and long-term social bonds, but whether it shapes the moment-to-moment dynamics of early social interactions is unclear. Here we examined how the intronic *Oxtr* single nucleotide polymorphism (SNP) NT213739 in female prairie voles (*Microtus ochrogaster*) shapes their dynamic social interactions with an opposite-sex conspecific. Leveraging a computational pipeline to analyze the movements of freely interacting voles, we found that C/C females, which expressed higher OXTR levels in the nucleus accumbens than T/T females, spent more time socially observing novel males from a distance, especially early in interactions. This genotype-phenotype relationship persisted in multiple contexts, including the social preference test. Thus, natural *Oxtr* variation biases social observation in females toward unfamiliar males before bonds form, consistent with models where accumbens OXTR enhances the salience of social cues. These findings show that SNPs can shape subtle behavioral dimensions in early social encounters, with important implications for the role of oxytocin in the study of social attachment.

## Introduction

The oxytocin receptor (OXTR) is a conserved G-protein-coupled receptor that regulates reproductive and social behaviors across species, including parturition [1], maternal care [2], mating [3], and pair bonding [4]. Genetic studies show that natural variation in the *Oxtr* gene predicts differences in brain OXTR expression, neural responses to social stimuli, social cognitive endophenotypes and long-term, coarse-grain measures of social affiliation in prairie voles and humans [4–7]. For example, an intronic *Oxtr* single nucleotide polymorphism (SNP NT213739) in male prairie voles strongly associates with social attachment differences and brain region-specific variation in OXTR levels [8], including in the NAc, which has been causally related to the emergence of a pair bond [9–11]. However, most of our understanding about the relationship between NAc OXTR density and pair bonding comes from the partner preference test (PPT) assay in voles [12], which is only one type of proxy for attachment that is often quantified by measuring side-by-side huddling with a cohabitated mating partner versus an opposite-sex stranger. In reality, pair bonding is a culmination of many iterated interactions that begin when individuals are first introduced and eventually lead to a preference to huddle with them, yet the role of the *Oxtr* genotype in shaping such finer-grained, dynamic social interactions is not yet known.

Studies exogenously manipulating the oxytocin system in a variety of species support the utility of finer grained behavioral analyses as a way to understand oxytocin’s role in modulating social behavior. For example, experiments delivering oxytocin intranasally to dogs, rhesus macaques, and humans show that it enhances transient features of social interactions such as social gaze [13–15]. Intranasal oxytocin also shifts social distance between conspecifics in a context-dependent manner [16–18]. Even in prairie voles, studies employing chronic administration of intranasal oxytocin have found effects on discrete categories of social behaviors, such as sniffing, lunging, investigation and contact [19–21]. Extracting dynamic social interactions in prairie voles to analyze their association with natural variation in the *Oxtr* gene should therefore yield new insights into oxytocin’s endogenous modulation of specific social interactions leading to a long-term attachment. Moreover, since *OXTR* SNPs in humans associate with neuropsychiatric phenotypes, a better understanding of the functional implications of *Oxtr*’s genetic variation could also positively impact the translation of personalized treatments to social deficits [22,23].

Advances in markerless, multi-individual, keypoint tracking of animals in video recordings [24–27] have set the stage for now analyzing dynamic social interactions in prairie voles with greater precision through time. Some have started to use these approaches [28], but they remain challenging to apply to freely interacting prairie voles, whose body shape and fur coloring make them difficult to separate. Nevertheless, as we show here, these tools can provide a detailed characterization of the relative body positioning between socially interacting prairie voles from the first moments of introduction through the first few hours of cohabitation and attachment formation. Thus, this type of quantification enables high-precision analyses of how oxytocin modulates prairie vole social interactions over time and across contexts.

Here we apply this approach to study the effects of natural variation in the *Oxtr* SNP NT213739 on prairie vole social interactions. Previous work showed that males homozygous for the C-allele have higher NAc OXTR expression and can form pair bonds more rapidly than males homozygous for the T-allele, suggesting that *Oxtr* genotype may effect early social interactions with a mating partner [8]. We now demonstrate, for the first time, that female prairie voles have the same genotype-linked difference in NAc OXTR levels, and our keypoint tracking analysis finds that this naturally occurring *Oxtr* SNP selectively biases stranger-directed social observation of the male before familiarity develops – a result that persists across social contexts.

## Material and Methods

### Animals

Adult (60 day+) male (n = 20) and female (n = 10 C/C and 10 T/T) prairie voles (*Microtus ochrogaster*) were used and originated from a laboratory breeding colony derived from field-captured voles in Champaign, Illinois. These animal numbers were chosen based on prior experience with pair bonding assays [29,30]. Animals were housed with a 14:10 h light/dark cycle at 68-72 °F and humidity 40-60% with *ad libitum* access to water and food (Laboratory Rabbit Diet HF # 5326, LabDiet; St. Louis, MO, USA). The cages contained Bed-o’Cobs Laboratory Animal Bedding (The Andersons, Maumee, Ohio), and environmental enrichment, which included cotton pieces to facilitate nest building. Animals were weaned at 20-23 days of age and group housed (2-3 per cage) with age- and sex-matched pups. All experiments were performed during the light cycle (between 9 a.m. and 5 p.m.). The females were not ovariectomized or estrogen-primed. All voles were sexually naïve and not pair-bonded at the time of the initial pairing for cohabitation, which lasted 48 hours – a duration not uncommonly used [31,32] and sufficiently long for pair bonds in (non-genotyped) wildtype voles to be established [33,34]. All experimental procedures were conducted in strict accordance with the guidelines established by the National Institutes of Health and were approved by the Emory University Institutional Animal Care and Use Committee.

### Animal breeding

Homozygous breeder pairs were used to produce T/T and C/C animals using previously described protocols (Boender et al., 2023). Female voles used for behavior recording were either T/T or C/C, and the male voles were all randomly selected from our outbred colony.

### Autoradiography

Fresh-frozen brains were sectioned on a cryostat (Epredia Cryostar NX-70, Thermo Fisher Scientific, MA, USA) at 20 μm, mounted on Superfrost Plus slides (Fisher Scientific), and stored at −80°C until use. Autoradiography was performed as previously described [30]. Briefly, slides were thawed and fixed for 2 min in 0.1% paraformaldehyde in phosphate-buffered saline (PBS) for 2 min, washed in 50 mM tris in PBS (pH 7.4, 2 × 10 min), and incubated in 50 mM tris buffer, supplemented with 0.1% bovine serum albumin and 50 pM I125-OVTA (2200 Ci/mmol, ornithine vasotocin analog, #NEX254010UC, PerkinElmer, MA, USA) at room temperature (RT) for 1 hour. Unbound ligand was removed by washing in 50 mM tris with 0.2% MgCl2 at 4°C (4 × 5 min) and 30 min at RT. Slides were dipped in Milli-Q, dried, and placed in a cassette with BioMax MR film (Sigma-Aldrich, MO, USA). After 7 days, films were developed and imaged using an MCID core system (Interfocus Co., UK). Mean gray values of the nucleus accumbens (NAc) were quantified in ImageJ. For densitometric calibration, the mean gray value of an unexposed area of the autoradiographic film was defined as the zero point in the calibration curve, such that all measurements were implicitly background-corrected during conversion to I¹²⁵ activity (disintegrations per minute) using the I¹²⁵ standard. NAc OXTR density levels of individual animals were determined by averaging values across at least five sections spanning the anterior-posterior axis of the NAc and compared between T/T and C/C animals.

### Experimental design

All recordings were conducted in a designated behavioral recording room that was separate from the animal colony. During each cohabitation recording, a T/T or C/C female and a WT male were placed in a 30 × 28 × 30 cm (length × width × height) rectangular arena with bedding, food, and water gel. The animals were allowed to freely interact and cohabitate. The first 165 min of cohabitation were recorded using a top-down Basler a2A1920-160ucPRO video camera at 30 frames per second. The animals were lightly shaved to assist with the ground truth data.

During each Social Preference Test (SPT) recording, each of the n = 20 T/T or C/C females was placed in a 75 × 20 × 30 cm rectangular arena with bedding. The partner and stranger males were each placed under rectangular metallic mesh containers (5.6 × 15 × 5.6 cm) at opposite ends of the arena. One body length (6.25 cm) around the container edges was defined as the social zone. Note that the SPT is distinct from the traditional PPT, which tethers the opposite sex conspecifics (partner and stranger) to different corners of an arena and allows otherwise free interactions with a nearby subject. We therefore refer to the SPT and social preferences when referencing results using mesh-contained voles as one proxy for pair bonding [29], and PPT when referencing studies using tethers – many of which were able to use the freer interaction to document other proxies of pair bonding, such as stranger-directed aggression [35].

### Keypoint Tracking and Pose Estimation

A customized markerless multi-animal pose estimation pipeline was used for animal tracking (https://github.com/senakoko/AutoPoseMapper). Raw video frames of interacting prairie voles were extracted using SLEAP/maDLC [24,25], and twenty-one body landmarks were labeled and trained on each animal. The trained networks were used to make predictions on new raw videos, and an autoencoder was applied to the SLEAP/maDLC tracked data. Raw videos were processed in parallel to SLEAP/maDLC using idtracker.ai to automate the generation of individual vole identities. This pipeline outputs key point tracking coordinates for each animal on a frame-by-frame basis. Instances of erroneous coordinates and swapped identities were curated using a customized graphical user interface (https://github.com/senakoko/poseCorrectionGUI) prior to behavioral analyses using MATLAB (R2024b). Manual curation was applied on a frame-by-frame basis for the first 15 min of cohabitation (27,000 frames per video) for all n = 20 subjects. After the first 15 min, curation was performed on a one-second basis. Two cohabitation recordings (both from T/T subjects) failed after the first 15 min and were dropped from analyses that spanned the full cohabitation.

### Machine Learning Features

Twenty-one body landmarks were extracted from each animal using 2D pose estimation. To enable the automated classification of prairie vole social behaviors, we derived 26 kinematic and geometric features following approaches similar to those described previously [36]. These features were computed on a frame-by-frame basis from the body landmarks sampled at 30 Hz and were designed to capture both within-animal dynamics and inter-animal spatial relationships. For improving classifier performance, several features were normalized and additionally processed using Gaussian smoothing. See **Supplemental Methods** for further details.

### Supervised behavior classification using random forest

To automate the classification of social behaviors, we trained random forest models using manually annotated video data. Frame-wise labels were obtained for three behaviors—approach, flee, and huddle—from the first ∼15 min of each video. An approach was defined as continuous forward movement toward the prospective mate’s center of mass within a relative heading angle (≤30°) initiated from ≥1 body length; flee was defined by rapid movement away from the other animal initiated from ≤2 body lengths; huddle was defined as sitting together with bodily contact. Attack and mounting/mating behaviors were not reliably found across pairs and so were not classified. In total, the annotated dataset comprised 363 approach episodes (2,003 frames), 153 flee episodes (1,205 frames), and 95 huddle episodes (35,665 frames).

We implemented and compared multiple classification algorithms using Python (version 3.11.4) and Scikit-learn (version 1.3.1) on a pilot dataset for testing feasibility and model selection. The Random Forest Classifier (implemented in scikit-learn [37]) achieved the best overall performance and was thus selected for final use. Each behavior-specific classifier was trained as a binary classifier. To address class imbalance (e.g., approach episodes being sparse), the majority class (non-behavior) was randomly under-sampled. The models were then trained with 5-fold cross-validation across the annotated segments of both animals from all 18 video clips (10 C/C females, 8 T/T females, and 18 WT males). For each model, the train–test split was performed at the bout level (continuous sequences of frames from a single behavioral episode, see below) rather than on individual frames. Approximately 80% of behavior bouts were randomly assigned to the training set, with the remaining ∼20% for testing. The trained classifiers achieved frame-wise (bout-wise) accuracies of 98% (100%) for approach, 97% (100%) for flee, and 91% (93%) for huddle behaviors. These performance levels are comparable to those reported in prior supervised behavior classification studies [36,38].

For analysis purposes, the frame-wise predictions were transformed into individual behavior episodes or bouts [36] with a two-step smoothing procedure. First, predicted behavior frames were smoothed by merging consecutive predictions separated by fewer than a predefined gap (e.g., 5 frames). Subsequently, any resulting behavior bouts shorter than a minimum duration threshold (e.g., 5 frames) were discarded to reduce false positives. This approach improved alignment with ground-truth behavior durations and was validated using bout-wise precision and recall.

### Generalized linear modeling of contact-dependent approach behavior

To investigate how genotype and social-spatial context influenced the expression of approach behavior, we fit a Generalized Linear Model (GLM) using R (version 4.3.1). Negative binomial distribution was used to account for overdispersion in behavioral count data. The dataset consisted of 15 min time blocks (n = 11 per recording) across 18 recordings (10 C/C and 8 T/T females; two T/T females were excluded for analyses due to video camera recording errors). The dependent variable was the number of predicted approach bouts in each 15 min time block. Independent variables included genotype (C/C or T/T), time block index, and spatial context captured using two continuous predictors—the average relative heading angle and the separation distance between the animals, normalized by average body length. Modeling was performed in R using the glm.nb function from the MASS package (version 7.3-60).

Model selection was carried out using Corrected Akaike Information Criterion (AICc). Candidate GLMs spanning three-way and four-way interaction scopes were compared using AICc-based greedy backward selection. This procedure did not remove any terms from the full four-way model. The full model achieved a lower AICc and a significant nested likelihood-ratio (χ²) test (p = 0.027) relative to the best reduced three-way alternative. Akaike weights were also computed, with the full four-way model accounting for ∼75% of the total Akaike weight, which was therefore used.

### Quantifying social observation behavior

Social observation behavior was heuristically defined as sitting far away (separation distance greater than two average body lengths of 12.5 cm) and observing (relative heading angle < 30°) without moving (center body and nose are each < 0.20 cm/s).

As a nonsocial control, we also considered observation of a static location in a cage, defined for each video based on the corner that was least occupied by the male vole through the full duration of the recorded cohabitation. This location was determined for each video separately by generating a heat map of the frame-by-frame location of the male’s center-of-mass, and binning to obtain an occupancy probability at a resolution of ∼1x1 cm. Summing these occupancies over the bins falling within two average body lengths of a corner, we then determined which corner had the lowest occupancy. Observation of this least-occupied corner was then heuristically defined as sitting far away (separation distance greater than two average body lengths) and observing (relative heading angle < 30°) that corner without moving (center body and nose are each < 0.20 cm/s).

### Statistical analysis

For paired comparisons of the autoradiography in **Fig 1B**, assumptions of normality and homogeneity of variance were assessed using the Shapiro–Wilk test and Levene’s test, and Mann–Whitney U test was used due to the nonparametric nature of the data. For analyses involving the repeated measures over time (**Figs 2F-I, 3B-D, 5B-C**), linear mixed-effects models (LME) were employed in MATLAB (fitlme), with subject included as a random effect. When residual diagnostics indicated deviations from model assumptions in cases such as heteroscedasticity, outcome variables were log-transformed prior to analysis, and model assumptions were re-evaluated (note in figure legends). We followed this with a post-hoc multiple comparisons-corrected test (multcompare using Tukey’s Honest Significant Difference) when warranted. For assessing main effects and interactions during the SPT (**Fig 1D, 6A-C**), two-way ANOVAs were used in MATLAB (anovan), similarly with log transformation applied when model assumptions were violated. Parallel analyses were also conducted with LMEs and did not alter our main conclusions around social observation (reported in **Supplemental Materials**). Figures display group means ± standard error of the mean (SEM) with individual animals plotted as points. Significance in figure panels are indicated by asterisks: *0.05, **0.01, ***0.001.

**Fig. 1.**
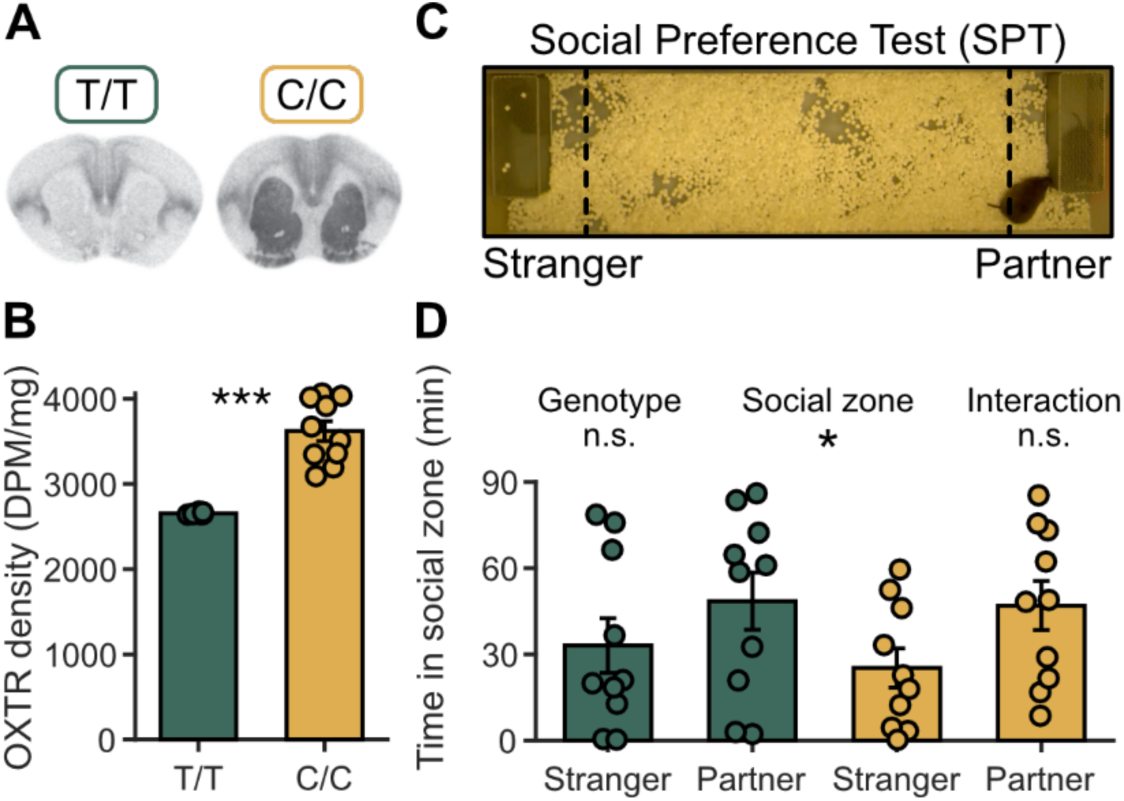
*Oxtr* genotype predicts NAc OXTR levels, while a social preference for the partner is intact across genotypes. (**A**) Representative autoradiographic images of OXTR binding in coronal sections from female prairie voles. (**B**) Quantified OXTR binding in nucleus accumbens (NAc) is higher in C/C than T/T females. Bars show mean ± SEM, and *** p<0.001, here and in subsequent figures, unless otherwise noted. (**C**) Social preference test apparatus; dashed lines indicate Stranger and Partner social zones. (**D**) In the SPT, females of both genotypes spent more time in the Partner than Stranger zone. * p<0.05, here and in subsequent figures, unless otherwise noted.

**Fig. 2.**
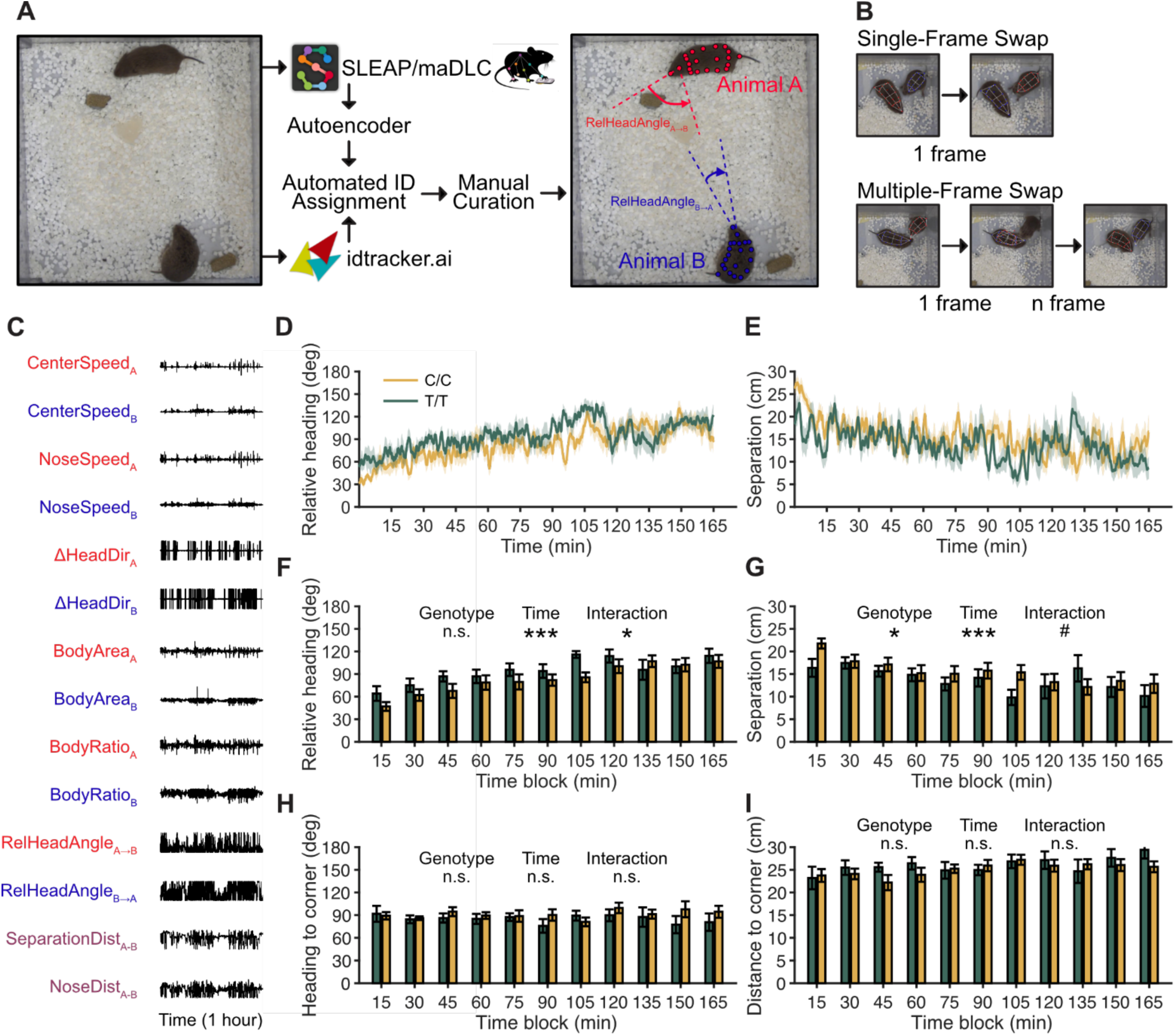
Markerless multi-animal pose estimation reveal an *Oxtr*-linked bias in social orientation during early cohabitation. (**A**) Novel pairs of female (T/T, *n* = 8, or C/C, n = 10 per group) and male wild-type (WT, *n* = 18) prairie voles interacted freely in an open field arena (30 x 28 cm), which was lined with bedding and provided *ad libitum* access to food and water gel. The first 165 min of a 48-hour cohabitation period were recorded from a top-down view using a camera capturing at 30 frames per second. A markerless multi-animal pose estimation pipeline (https://github.com/senakoko/AutoPoseMapper) was used to track the voles’ body part key points and movements. (**B**) Common pose tracking errors included single-frame swaps (identities of the two prairie voles were swapped for one frame) and multiple-frame swaps (key points for both voles were incorrectly assigned to a single subject over multiple frames before correcting back to two subjects). (**C**) From curated body coordinates we computed frame-wise features of both the female (red) and male (blue), including relative heading angle (female head direction vs. vector to male; 0° = facing male) and separation distance (centroid-centroid). (**D**) Relative heading angle changed over time during the first 3 h of cohabitation. (**E**) Separation distance also changed over time during the first 3 h. (**F**) In 15 min bins, C/C females oriented more directly toward the male than T/T. ** p<0.01, here and in subsequent figures, unless otherwise noted. (**G**) Separation distance showed a trend toward a genotype effect across 15 min bins. # 0.05<p<0.1, here and in subsequent figures, unless otherwise noted. (**H-I**) Nonsocial controls showed no genotype or time effects: Heading toward the arena center (H) and distance to center (I) were stable (all p > 0.10), indicating that genotype differences in (F-G) reflect social rather than spatial habituation factors.

## Results

The intronic single nucleotide polymorphism at NT213739 in the oxytocin receptor (OXTR) gene robustly predicts OXTR expression levels in the nucleus accumbens (NAc) and other brain areas of male prairie voles [8]. Here we quantified NAc OXTR expression in female prairie voles (n=10 per genotype) and observed (**Fig 1A-B**) the same result (Mann-Whitney *U*, *U* = 0.00, z = -3.74, p < 0.001).

Since NAc OXTR levels are implicated in long-term attachments [3,8,39], we next assessed a proxy of such pair bonding using a social preference test (SPT, **Fig 1C**). We cohabitated both high-expressing C/C and low-expressing T/T females with partner males for 48 hours. We chose not to ovariectomize or hormonally prime the females to allow the social behaviors to occur as naturally as possible, especially since endogenous estrogens can interact with oxytocin signaling [40]. In the SPT (**Fig 1D**), we found a significant main effect of social zone (Partner vs. Stranger), but no genotype or interaction effects (two-way ANOVA: main effect of genotype, F(1,36) = 0.28, p = 0.598, partial η² = 0.007; main effect of social zone, F(1,36) = 4.47, *p* = 0.041, partial η² = 0.099; social zone x genotype interaction, F(1,36) = 0.13, p = 0.724, partial η² = 0.003). Hence, on this long cohabitation timescale, females of both genotypes form social preferences for the partner despite a vast difference in NAc OXTR levels – seemingly indicating that the OXTR is not the only determinant of whether a partner preference can emerge, as also suggested by a recent OXTR knock-out study [41].

The absence of a genotype difference in the long time-scale preference for a partner does not preclude the possibility that females of these two genotypes differ in the dynamics of social interactions leading up to a pair bond. Indeed, previous work shows that C/C males, unlike T/T males, are able to form partner preferences after an abbreviated (6-hour) cohabitation, suggesting that *Oxtr* genotype may affect the early social interactions between mating partners. Since wild type female prairie voles can form attachments more quickly than males [33,34], we examined free interactions by C/C and T/T females during the first ∼3 hours of cohabitation when their cohabitated male was more of a stranger. We utilized a computational pipeline combining several machine learning algorithms (see **Methods**) to accurately track keypoints on the bodies of socially housed individuals in an unbiased fashion and estimate their poses (**Fig 2A**). As powerful as these algorithms now are, animal swap errors were not uncommon (**Fig 2B**) and were detrimental for social analyses requiring correct assignment of behaviors to individuals. Thus, we added a final manual curation step (see **Methods**) before generating behavioral features as a function of time (**Fig 2C**).

By tracking social behavioral features frame-by-frame, we observed that the relative heading angle toward the prospective mate (**Fig 2D**) and the separation distance between individuals (**Fig 2E**) changed over the course of the first three hours of cohabitation. Quantifying this in 15 min time blocks, both genotypes spent significantly more time initially facing toward the prospective mate (<90°, **Fig 2F**) and farther apart (**Fig 2G**). Moreover, we saw a significant interaction between genotype and time on the relative heading angle (LME, main effect of Genotype: F(1,176) = 2.20, p = 0.14, partial η² = 0.012; Time: F(10,176) = 14.183, p = 3.03x10^-18^, partial η² = 0.446; and interaction, F(10,176) = 2.00, p = 0.036, partial η² = 0.102). Post-hoc analysis of the interaction indicated that the relative heading angles for C/C females early in the cohabitation (up to 45 min) were typically significantly lower than that for T/T females later in the cohabitation (>75 min), while no such differences were seen for T/T females early on compared to C/C females later (multiple comparison t-test with Tukey’s HSD, p < 0.05).

At the same time, we observed a significant main effect of genotype on the separation distance (LME, main effect of Genotype: F(1,176) = 4.821, p = 0.029, partial η² = 0.027; Time: F(10,176) = 4.098, p = 4.35x10^-5^, partial η² = 0.189; and interaction, F(10,176) = 1.644, p = 0.098, partial η² = 0.085). Hence, *Oxtr* genotype was associated with how animals positioned themselves relative to each other over the course of social interaction, with C/C females typically oriented more than T/T females towards the male early on, while staying further away.

To check whether this phenotypic difference was specific for orienting towards a social versus nonsocial location, we next analyzed the same positioning measures relative to a static corner in the cage. Since rodents tend to crowd along the edges of an arena, we generated an occupancy map (see **Methods**) of a male vole’s location throughout a recorded cohabitation and chose for each video the least-occupied corner as a nonsocial location the female oriented towards with minimal contamination by the presence of the male. We found no significant effects of genotype, time, or any interaction on the relative positioning to this corner. Specifically, the average heading angle toward the male vole’s least-occupied corner **(Fig 2H)** was largely stable (LME, main effect of Genotype: F(1,176) = 0.067, p = 0.797, partial η² = 3.78x10^-4^; Time: F(10,176) = 0.758, p = 0.669, partial η² = 0.041; and interaction, F(10,176) = 0.823, p = 0.607, partial η² = 0.045). This was also true of the distance to the male vole’s least-occupied corner **(Fig 2I)** (LME, main effect of Genotype: F(1,176) = 0.073, p = 0.788, partial η² = 4.13x10^-4^; Time: F(10,176) = 1.370, p = 0.198, partial η² = 0.072; and interaction, F(10,176) = 0.939, p = 0.499, partial η² = 0.051). Thus, the differences over time and genotype in a pair’s relative positioning rather than just habituation to the spatial structure of the enclosure. Together, these results motivated us to look next at specific social interactions occurring during this early cohabitation phase.

We used the keypoint data to train a supervised machine learning algorithm (Random Forest Classifier, see **Methods**, **Fig 3A**) to predict the occurrence of three particular social interactions that all animals displayed: individual flee (**Fig 3B**), mutual huddle (**Fig 3C**), and individual approach (**Fig 3D**). We did not observe a significant main effect of, or interaction between, genotype or time on the number or duration (**Fig 3B**) of flee events (LME, raw data log transformed, main effect of Genotype: F(1,176) = 1.155, p = 0.284, partial η² = 0.007; Time: F(10,176) = 3.120, p = 0.001, partial η² = 0.151; and interaction, F(10,176) = 1.065, p = 0.392, partial η² = 0.057). In contrast, huddling was expressed significantly more over time, as evidenced by the increasing duration of huddling epochs. There was no genotype main effect and only a trending interaction with time (LME, main effect of Genotype: F(1,176) = 0.173, p = 0.678, partial η² = 0.001; Time: F(10,176) = 3.757, p = 1.35x10^-4^, partial η² = 0.176; and interaction, F(10,176) = 1.862, p = 0.053, partial η² = 0.096), suggesting that both C/C and T/T females expressed their pair bond at similar rates during cohabitation.

**Fig. 3.**
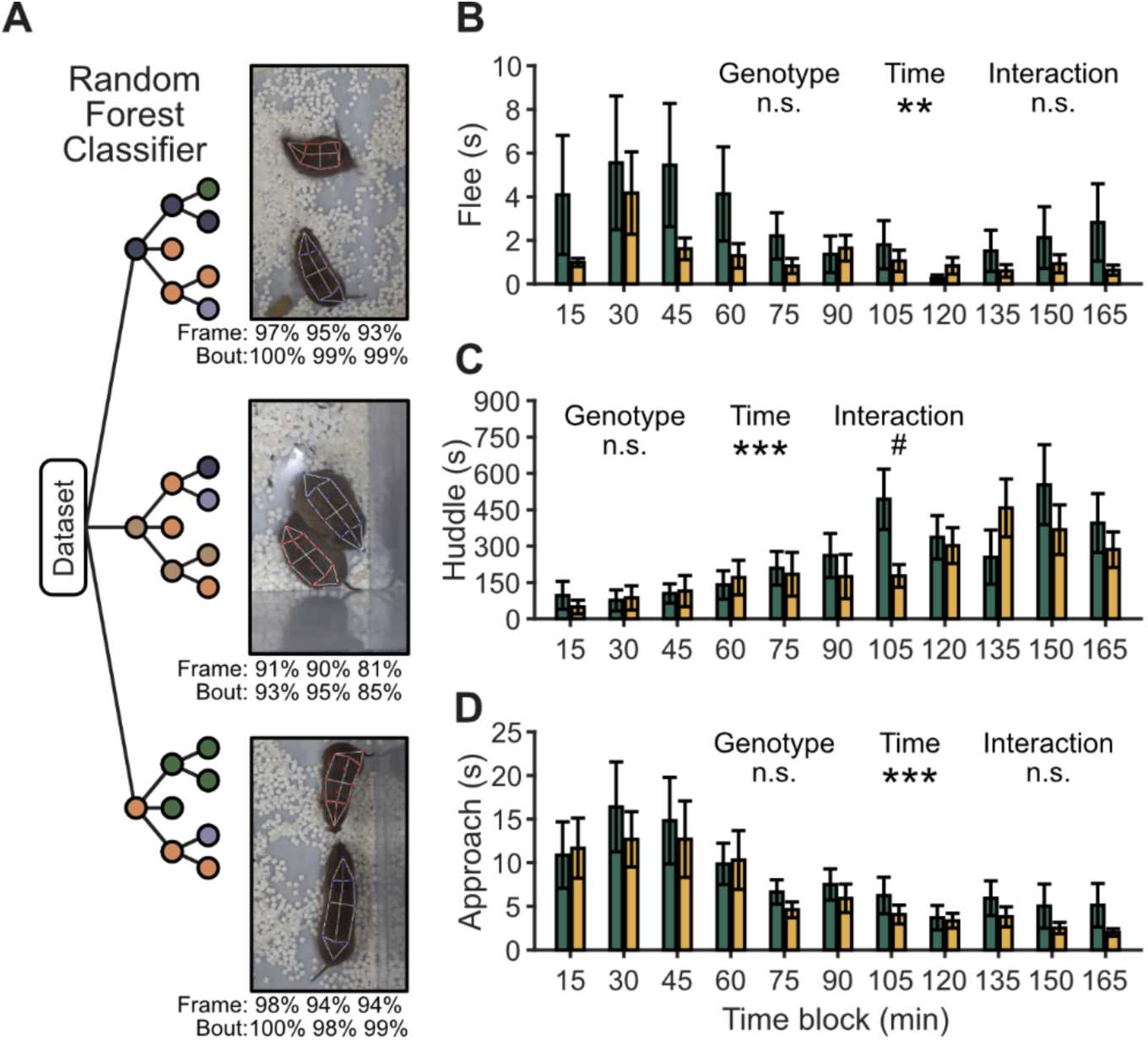
Automated supervised machine learning classification of social behaviors across time. (**A**) Spatiotemporal features from the pose tracking were used to train a random forest classifier to detect flee, huddle, and approach behaviors across the cohabitation. Classifier performance as reported by scikit-learn (Python) at the Frame (top) and Bout (bottom) levels are listed below each example frame, in order of Accuracy, Precision and Recall (left to right). (**B**) Flee behavior was transient and did not vary systematically across the cohabitation. Raw data log transformed prior to analysis. (**C**) Mutual huddle behavior increased over time for both genotypes. (**D**) Approach behavior decreased over time for both genotypes. Raw data log transformed prior to analysis.

We also saw a significant time dependence where the amount of time spent approaching was greater for both genotypes early in the cohabitation and decreased after the first hour – consistent with the idea that rodents investigate more a conspecific who is novel. We again saw no genotype main effect or interaction (LME, raw data log transformed, main effect of Genotype: F(1,176) = 0.012, p = 0.911, partial η² = 7.05 x10^-5^; Time: F(10,176) = 3.744, p = 1.40x10^-4^, partial η² = 0.175; and interaction, F(10,176) = 0.130, p = 0.999, partial η² = 0.007).

The lack of a genotype effect for the time spent approaching was unexpected, given indications that the relative heading angle (**Fig 2F**) and separation (**Fig 2G**) were at least somewhat modulated by genotype, and the fact that social approach involves directed movement toward the other animal. This led us to ask whether the likelihood of an approach event occurring, rather than the duration of approach, might be predicted by these factors in a genotype-dependent manner. We applied a generalized linear model (GLM) that included relative heading angle, separation, genotype and the 15 min time blocks as factors to predict the occurrence of approach events (**Table 1**).

**Table 1.**
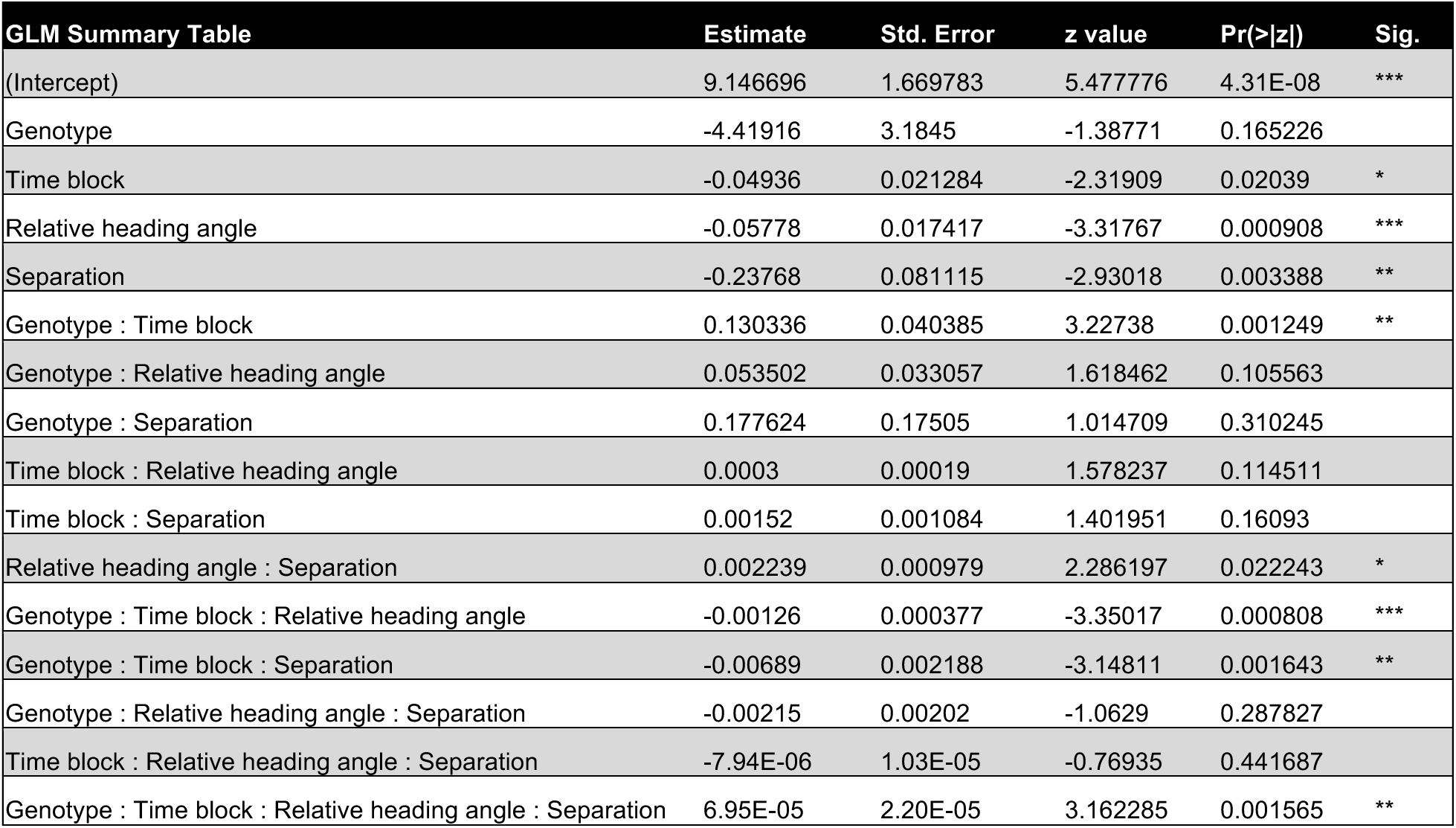
GLM to predict the rate of social approach events by genotype, time, and keypoint kinematics.

| GLM Summary Table | Estimate | Std. Error | z value | Pr(>\|z\|) | Sig. |
| --- | --- | --- | --- | --- | --- |
| (Intercept) | 9.146696 | 1.669783 | 5.477776 | 4.31E-08 | *** |
| Genotype | -4.41916 | 3.1845 | -1.38771 | 0.165226 |  |
| Time block | -0.04936 | 0.021284 | -2.31909 | 0.02039 | * |
| Relative heading angle | -0.05778 | 0.017417 | -3.31767 | 0.000908 | *** |
| Separation | -0.23768 | 0.081115 | -2.93018 | 0.003388 | ** |
| Genotype : Time block | 0.130336 | 0.040385 | 3.22738 | 0.001249 | ** |
| Genotype : Relative heading angle | 0.053502 | 0.033057 | 1.618462 | 0.105563 |  |
| Genotype : Separation | 0.177624 | 0.17505 | 1.014709 | 0.310245 |  |
| Time block : Relative heading angle | 0.0003 | 0.00019 | 1.578237 | 0.114511 |  |
| Time block : Separation | 0.00152 | 0.001084 | 1.401951 | 0.16093 |  |
| Relative heading angle : Separation | 0.002239 | 0.000979 | 2.286197 | 0.022243 | * |
| Genotype : Time block : Relative heading angle | -0.00126 | 0.000377 | -3.35017 | 0.000808 | *** |
| Genotype : Time block : Separation | -0.00689 | 0.002188 | -3.14811 | 0.001643 | ** |
| Genotype : Relative heading angle : Separation | -0.00215 | 0.00202 | -1.0629 | 0.287827 |  |
| Time block : Relative heading angle : Separation | -7.94E-06 | 1.03E-05 | -0.76935 | 0.441687 |  |
| Genotype : Time block : Relative heading angle : Separation | 6.95E-05 | 2.20E-05 | 3.162285 | 0.001565 | ** |

Both relative heading angle and separation were significant predictors by themselves, as expected for this kind of behavior. Moreover, time block was also significant, consistent with the time dependence of approach during early cohabitation (**Fig 3D**). Importantly, several interactions were significant, including those between genotype, time block and either relative heading angle (p<0.001) or separation (p<0.005) or both (p<0.005). In other words, C/C and T/T animals differed across time in how their relative heading angle or separation during a time block predicted whether they approached the other individual.

To better visualize the interplay between these variables, we generated the genotype-specific joint distributions of separation distance and relative heading angle for all frames in each time block (**Fig 4A**). Screenshots from different vole pairs illustrate relative body positions corresponding to various combinations of these variables in the joint distribution. Hotspots in the distributions reflect oft-occurring positions, such as when a pair huddled at close distance and faced opposite directions from each other.

**Fig. 4.**
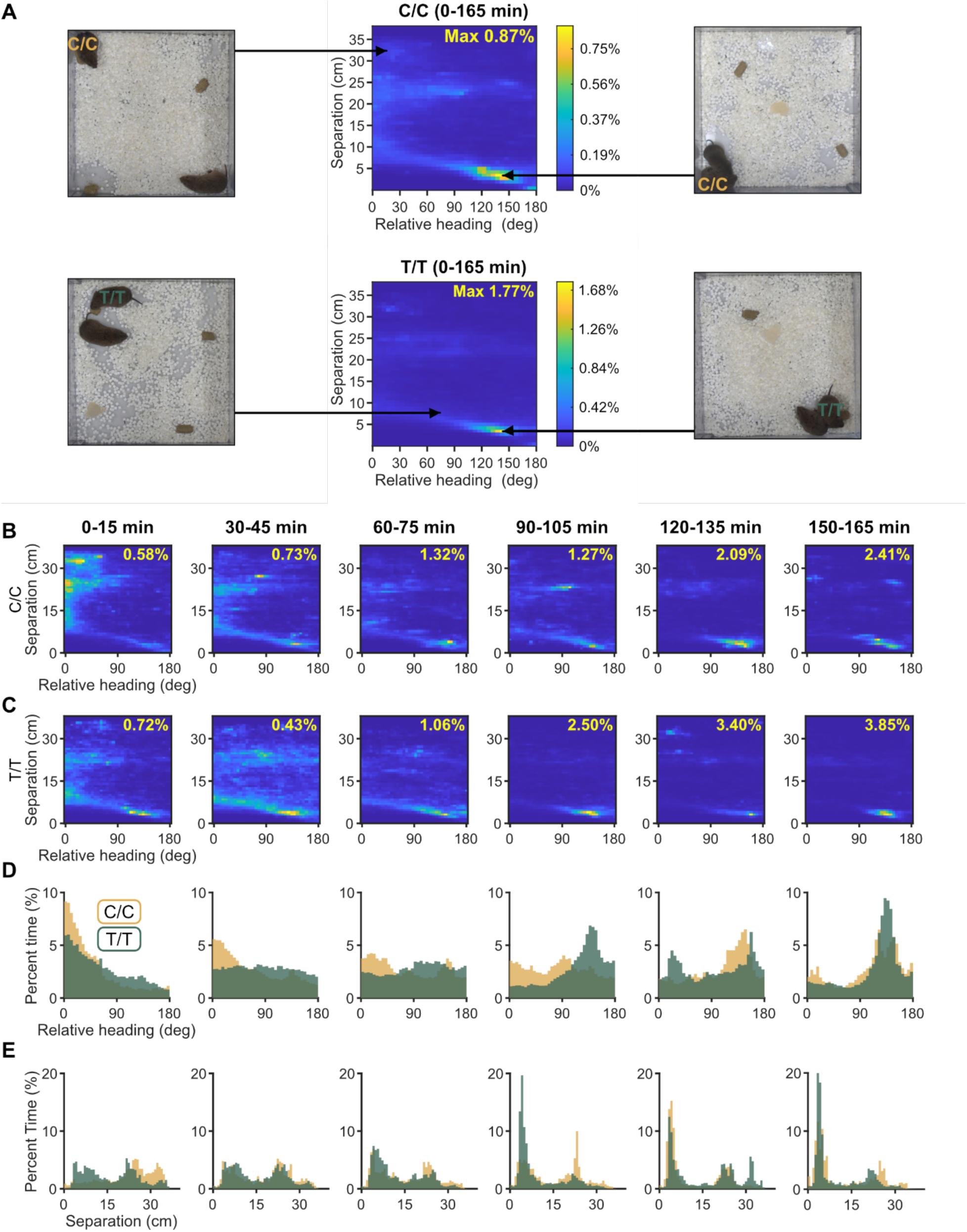
Joint orientation-distance kinematics reveal time- and genotype-dependent social configurations. (**A**) Genotype-specific joint distributions (2D occupancy maps) of relative heading angle (female head direction vs. vector to the male centroid; 0° = facing male) and separation distance (centroid-centroid) computed from all frames across the full 165 min cohabitation. Max percent time across all bins is shown as yellow text within each map. Insets show example frames illustrating frequent configurations (e.g., close distance with opposite facing bodies during huddling). (**B-C**) Joint distributions binned in 15-min intervals for C/C (**B**) and T/T (**C**) females reveal distinct early-cohabitation configurations that converge over time. (**D**) Proportion of time oriented toward the male in each 15-min bin: C/C females show greater early orienting than T/T, with differences most pronounced in the first 15 min and diminishing thereafter. (**E**) Proportion of time for separation distance between the female and male: C/C females are farther from the male in the first 15 min, whereas both genotypes spend most time at close distances by the final bin.

Over the course of the entire 165 min video, the distributions for C/C and T/T females appeared qualitatively similar (**Fig 4A, top vs. bottom heat maps**). However, broken down into 15 min time blocks, C/C and TT females positioned themselves differently relative to their male in a stereotyped fashion (**Fig 4B-C**). This was especially obvious in the first 15 min when C/C females spent more time oriented toward (<30°, **Fig 4D, leftmost panel**) and farther away from (**Fig 4E, leftmost panel**) the male. By the last 15 min (**Fig 4DE, rightmost panel**), both C/C and T/T animals spent most of their time next to each other, as expected given the time course of huddling behavior (**Fig 3C**).

The conjunction of a narrower relative heading while being farther apart is suggestive of a construct such as “social observation,” where animals are heightened in their oriented watchfulness of another individual. We checked for this behavior directly by defining a heuristic on the keypoints (see **Methods**) to automatically extract epochs when C/C and T/T females were *motionlessly* observing the male from a distance of at least two body lengths (**Fig 5A**). Both C/C and T/T animals exhibited such observation behavior (**Fig 5B**). However, we found a significant main effect of genotype and time block on observation duration, and a significant interaction between the two (LME, main effect of Genotype: F(1,176) = 14.06, p = 2.40x10^-4^, partial η² = 0.074; Time: F(10,176) = 10.587, p = 6.01x10^-14^, partial η² = 0.376; and interaction, F(10,176) = 2.866, p = 0.0025, partial η² = 0.140). Post-hoc analysis of the interaction indicated that social observation by C/C females in the first 15 minutes was significantly higher than that for T/T (and C/C) females later in the cohabitation (>60 min), while no such differences were seen for T/T females (multiple comparison t-test with Tukey’s HSD, p < 0.05). In fact, consistent with the joint distributions during the first 15 minutes of the cohabitation (**Fig 4B-C**), C/C animals spent more than twice the time compared to T/T animals observing the then-novel male.

**Fig. 5.**
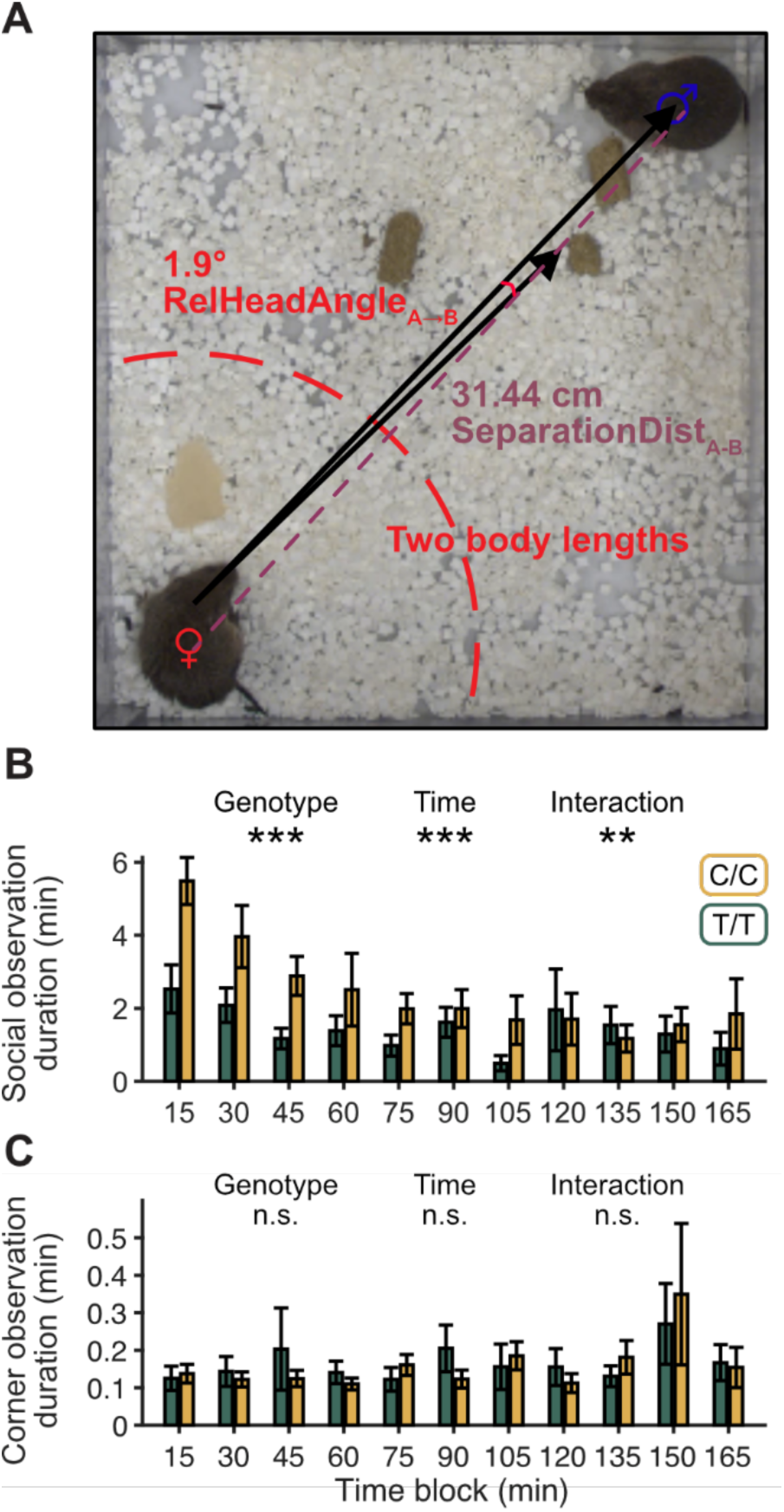
Heuristic detection of social observation reveals an *Oxtr*-dependent, temporally evolving phenotype. (**A**) From curated pose tracking, “social observation” epochs were automatically identified when the female was stationary, oriented toward the male (Relative heading angle < 30°, and at a centroid-centroid separation distance > two body lengths (dashed curve). (**B**) Quantification of the female’s social observation duration toward a novel male in 15-min bins. (**C**) Quantification of the female’s observation duration toward the male’s least-occupied corner in 15-min bins. Raw data log transformed prior to analysis.

By comparison, observation towards a nonsocial, static location (the corner least occupied by the male vole) was not significantly affected by genotype, time, or any interaction between the two **(Fig 5C)** (LME, raw data log transformed, main effect of Genotype: F(1,176) = 0.038, p = 0.845, partial η² = 2.18x10^-4^; Time: F(10,176) = 0.499, p = 0.889, partial η² = 0.028; and interaction, F(10,176) = 0.344, p = 0.968, partial η² = 0.019). Thus, the differences over time and genotype in the duration of the female’s observation of the male were likely specifically attributable to social factors.

The intriguing possibility that C/C females show a distinctive social phenotype toward novel individuals led us finally to examine whether such observation behavior would also manifest in a genotype-dependent manner in a different context, such as for a Stranger male in the SPT. We analyzed the first 15 min of the SPT when the Stranger was novel and compared that to the first 15 min of the cohabitation when the soon-to-be Partner was also novel. In the first 15 min of the SPT, we found a significant interaction (but no main effects) in the amount of time that C/C versus T/T females spent near the Partner compared to the Stranger (**Fig 6A**, two-way ANOVA, main effect of genotype: F(1,36) = 1.46e-04, p = 0.990, partial η² = 3.50e-06; social zone: F(1,36) = 0.05, p = 0.826, partial η² = 0.001; and interaction, F(1,36) = 5.64, p = 0.02, partial η² = 0.119). Thus, after a 48-hour cohabitation, C/C and T/T females tended toward opposite preferences in which conspecific they socially investigated when both the Partner and a Stranger were present.

**Fig. 6.**
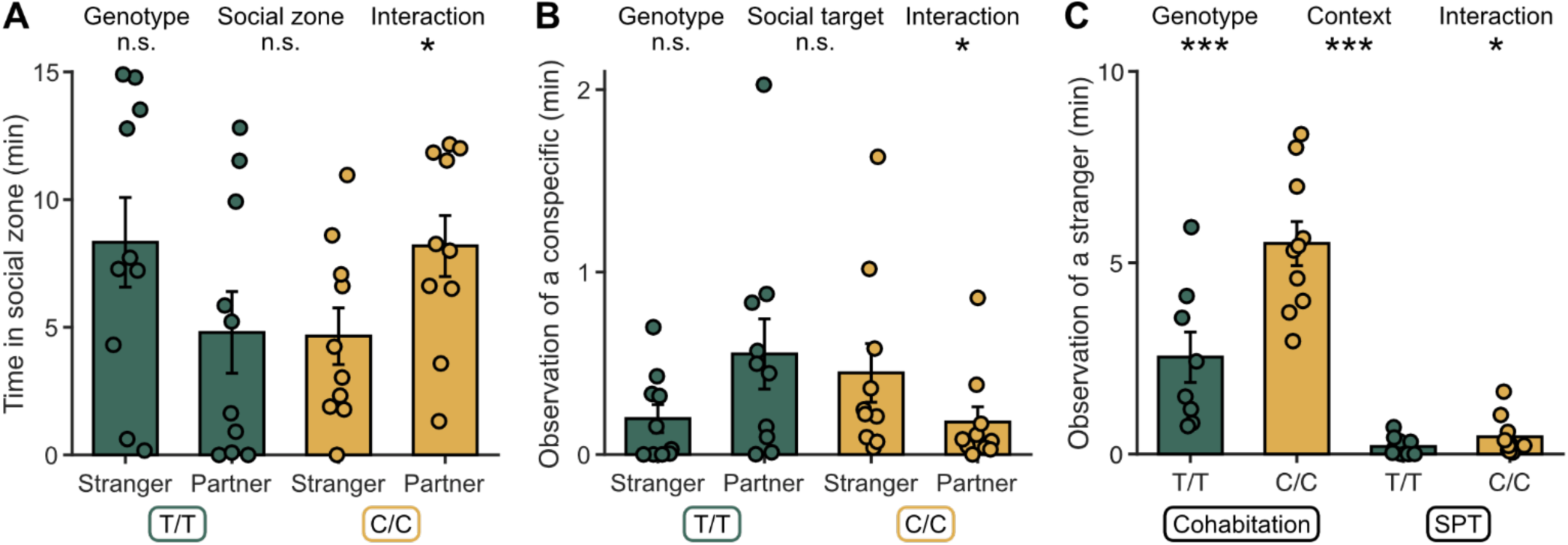
Context dependence of *Oxtr*-linked social observation of a stranger. (**A**) During the first 15 min of the SPT, females showed a Genotype × Social zone interaction in time spent near Stranger vs. Partner. Average T/T females spent more time with the stranger, whereas C/C spent less time. (**B**) During the first 15 min of the SPT, females showed a Genotype × Social target interaction in time spent observing the Stranger vs. Partner. Average T/T females spent more time observing the Partner, whereas C/C more time observing the Stranger. Raw data log transformed prior to analysis. (**C**) Comparing social observation of a novel prospective mate during the first 15 min of cohabitation (novel male freely interacting) to the SPT (novel Stranger behind a barrier), C/C females consistently observe for longer than T/T females even though the SPT dampened overall observation. Raw data log transformed prior to analysis.

Intriguingly, we also saw a significant interaction (but no main effects) in the duration of social observation of the Partner versus the Stranger across genotypes (**Fig 6B**, two-way ANOVA, raw data log transformed, main effect of genotype: F(1,36) = 0.1847, p = 0.670, partial η² = 0.004; social target: F(1,36) = 0.061, p = 0.807, partial η² = 0.001; and interaction, F(1,36) = 5.62, p = 0.023, partial η² = 0.5). Mirroring the cohabitation, C/C females tended to observe the Stranger more in the SPT, while T/T females observed them less.

In fact, when comparing across cohabitation and SPT contexts, we found significant main effects of genotype and context on observation duration, as well as a significant interaction (**Fig 6C**, two-way ANOVA, raw data log transformed, main effect of genotype: F(1,34) = 15.737, p = 3.56x10^-4^, partial η² = 0.074; context: F(1,34) = 133.643, p = 2.51x10^-13^, partial η² = 0.403; and interaction, F(1,34) = 6.032, p = 0.019, partial η² = 0.030). Thus, while observation was overall muted in the SPT, C/C females nevertheless observed a novel male longer than T/T females across contexts.

## Discussion

Using a quantitative pipeline to discover subtle social behavioral differences [24,25,42], we found that the *Oxtr* SNP NT213739 in female prairie voles is associated with the degree of observation of a novel male in a social context-dependent manner. These are the first data to show that NT213739, which has previously been associated with neural activity patterns and partner preference formation in male prairie voles [5,8], is associated with social behaviors in females. By analyzing female behavior toward a novel male, we revealed a significant genotype-associated difference in the “style” of their social interactions. Females with the C/C genotype, which correlates with high levels of OXTR expression in the NAc, socially observed novel males more than T/T females, which expressed OXTR at lower levels in the NAc. A main effect of genotype held across behavioral contexts where the novel male was either freely moving or restricted from interacting with the female. Overall, our results suggest that *Oxtr* genotype is associated with differences in female behavior towards unknown males during early social interactions.

Previous studies of prairie vole social behavior have relied heavily on the PPT, where the key behavioral measure is time spent huddling with a tethered mating partner relative to an opposite-sex stranger over the course of several hours. In contrast, our study leveraged advances in computer vision and keypoint tracking to quantify more subtle dimensions of behavior through time. Such methods have grown considerably over the past decade [45] and have even been applied to long-term video recordings of prairie voles housed in separate compartments [28]. Building on these studies, we added an autoencoder architecture on top of the existing tracking software to improve standard pose estimation results during free interactions but found it still necessary to manually correct animal-swap errors. Compared to traditional manual scoring, two strengths of this approach are 1) sidestepping the effort required for one or more human observers to annotate many long videos, and 2) enabling measurement of subtle differences in behavior that would be difficult and/or impractical for human observers to analyze. Using this method, clear genotype-linked differences in social behavioral dynamics across multiple contexts emerged.

A common element across contexts was the fact that a genotype difference only appeared early on, suggesting that this *Oxtr* SNP may specifically modulate social interactions with a stranger, without affecting the ability to eventually show a social preference for the partner after a sufficiently long cohabitation. This finding about natural variation in the *Oxtr* gene impacting stranger interactions seems reconcilable with recent findings that exogenously knocking out this gene from birth in prairie voles does not impair the ability to form pair [41] or same-sex peer social relationships [46], given enough time. Over long cohabitation, the relative lack of OXTR – whether due to the T/T genotype or exogenous knock-out – does not apparently impede bond formation. One hypothesis is that OXTR impacts what actions are taken during the early stages of a social relationship, rather than just the overall likelihood of social bonding per se. Indeed, injecting females with oxytocin intracerebroventricularly [47] or overexpressing OXTR in their NAc [30] can drive a partner preference despite only short cohabitations; and *male* C/C voles, in contrast to T/T voles, form partner preferences within only a short 6 hour cohabitation [8]. Our results here suggest that one such early social action influenced by the *Oxtr* gene could be stranger observation, which captures the tendency to observe an unknown individual from afar.

We heuristically defined “social observation” from the time series of tracked keypoints in a manner similar to the “social vigilance” or “risk assessment” constructs often studied in the context of stress and threat [48–51]. Similarities included the requirement that one’s head be oriented toward the other’s location while outside of a direct interaction zone. However, we also required the observing animal to be stationary (defined by minimal frame-by-frame change in the body center-of-mass and head locations) and not simply momentarily oriented toward the other individual. This helped quantitatively discriminate social observation from social approach during naturalistic, unrestricted interactions between individuals. Finally, although observation and vigilance were operationally similar, the former term is broader and can apply whether animals are stressed or not.

Our “social observation” construct may be akin to “social gaze” – a term often applied to watching another individual, looking at their eyes, and following their gaze [52–54]. However, we did not characterize the behavior of the observed prairie vole or the contingent actions taken by the observer. Thus, we acknowledge that the two may not be strictly equivalent. Nevertheless, social gaze is a construct widely studied across the animal kingdom and is known to be modulated by exogenous oxytocin manipulations. In domestic dogs [15] and cats [55], experimentally elevating oxytocin levels is linked to longer owner-directed gazing. In non-human primates, oxytocin increases eye fixation [14] and gaze-following [56] in rhesus macaques. Human studies have similarly shown that intranasal oxytocin increases gaze toward the eye region of social stimuli [13,57,58]. To the degree that intranasal oxytocin increases oxytocin levels centrally [59,60], these studies across species suggest that central OXTR activation generally heightens social gazing behavior. Our finding that high OXTR females observed novel males more is consistent with this hypothesis and extends the idea to rodents.

Studies that exogenously manipulate the oxytocin system in mammals also support its role in modulating social distance, although in a context-specific manner [61]. Domestic dogs maintain a closer proximity to their familiar owners when given intranasal oxytocin [15]. Administering the same treatment to African lions decreases their distance to their nearest familiar neighbor in the presence of stranger cues [17]. In non-human primates, however, oxytocin increases social distance between familiar capuchin monkeys when they compete for food [16]. Meanwhile, in pair-bonded female marmosets, intranasal oxytocin biases animals toward closer social interactions with their partner rather than a stranger [62]. Finally, in humans, intranasal oxytocin can either decrease the ideal distance to an unknown attractive opposite-sex individual in women [63] or increase it in men with romantic partners [18].

In prairie voles, rather than employing exogenous manipulations of the oxytocin system [11,64,65] or *Oxtr* gene [29,41,66], we took advantage of natural genetic variation to demonstrate how temporal dynamics in a social context affect whether an endogenous *Oxtr* SNP impacts separation distance and social observation. Although our study design was limited in not including heterozygous females to assess intermediate phenotypes, we nevertheless found a main effect of genotype only early on when the male was novel, both when animals were freely interacting or restricted in the SPT. Intriguingly, this result is reminiscent of a similar correlation in humans between endogenous OXTR expression and gazing at an unfamiliar individual. Specifically, women carrying an A-allele in the *OXTR* SNP rs53576 gazed longer than GG-allele females at a novel interaction partner, but only in the first 5 min of their chat [67]. This SNP has been widely investigated for its role in human social behavior [68], and carriers of the A-allele generally have higher OXTR levels, as measured in several brain regions in post-mortem tissue [69]. Intriguingly, the same SNP predicts greater anxiety and stress reactivity in human [70,71], suggesting that C/C prairie voles could provide a translationally relevant model to investigate mechanisms of stress [20]. Indeed, the fact that our C/C voles showed both high OXTR in a brain region-specific fashion [8] and feature longer social observation times of novel conspecifics opens the door to using voles to better delineate how OXTR expression in particular brain areas facilitates social gaze-like phenotypes.

In fact, animal studies are already elucidating how specific limbic structures underlie the effects of stress on the “social vigilance” construct. Studies in California mice (*Peromyscus californicus*) have shown that females are more socially vigilant toward a novel same-sex conspecific when stressed, and antagonizing OXTR in the bed nucleus of the stria terminalis (BNST) weakens this vigilance [49,50]. The role that NAc OXTRs play in these females emerges under stress, since OXTR-Gq agonism reduces social vigilance in stressed females, but antagonism in unstressed females has no effect on social vigilance [51]. These results highlight how OXTRs in distinct brain areas mediate opposing effects on social vigilance, which may be important to factor in for future research given NT213739’s impact on OXTR expression in other brain areas [72].

## Conclusions

Our findings demonstrate that a naturally occurring intronic polymorphism in female prairie voles’ *Oxtr* gene influences their social observation of strangers. This phenotype emerged consistently across contexts yet did not impair the eventual formation of pair bonds. This result suggests that endogenous *Oxtr* variation shapes the details of social actions, rather than their long-term outcome. This finding is in agreement with the hypothesis that oxytocin heightens the salience of social cues [73–77], and has implications for real-world behavior. In nature, social encounters are often fleeting, meaning that these subtle differences in perceptual salience and attentional focus toward potential mates due to genetic differences in *Oxtr* could bias behavioral trajectories toward or away from pair bonding in less contrived contexts than studied in the lab. Thus, rather than focusing on *whether* OXTR mediates pair bonding, future inquiries should determine *how* OXTR modulates specific, potentially ephemeral, dimensions of social interactions, ideally in identified cell populations and circuits. The computational behavioral analysis tools employed here will facilitate such studies and can be combined with cell type and circuit-specific manipulations of OXTR and physiology to elucidate the oxytocin system’s specific effects on dynamic social interactions. By integrating genetic, neurochemical, and computational behavioral approaches, our study describes how natural variation can tune early social dynamics, opening new opportunities to dissect oxytocin’s role in mediating affiliative behaviors.

## Acknowledgements

This work could not have been done without the generous collaborative interactions and support from L.J.Y., who passed away before this work was completed. We dedicate this study to his memory, especially given his many contributions to the social neuroscience field and the idea that oxytocin modulates the salience of social cues. This work was funded by NIH grants R01MH115831 (G.J.B. and R.C.L.), P50MH100023 (L.J.Y. and R.C.L.), R35GM155357 (Z.V.J.), P51OD11132 (to Emory National Primate Research Center) and NIH-IRACDA Postdoctoral Fellowship K12GM000680 (S.L.T.L.). We thank Lorra Julian, Dr. Denyse Levesque, and the rest of the EPC animal care and veterinary staff for vole husbandry and care.

## Author Contributions (CRediT) Statement

Conceptualization: S.A., G.J.B., L.J.Y., R.C.L.

Methodology: S.A., X.M.C.

Software: S.A., S.L.T.L., X.M.C.

Investigation: S.A., S.L.T.L., X.M.C.

Data curation: S.A., S.L.T.L., X.M.C., C.B.

Formal analysis: S.L.T.L., X.M.C.

Funding acquisition: G.J.B., L.J.Y., R.C.L.

Project administration: R.C.L.

Resources: A.J.B.

Supervision: G.B.J., L.J.Y., R.C.L.

Visualization: S.L.T.L.

Writing-original draft: S.L.T.L., X.M.C., Z.V.J., R.C.L.

Writing-review & editing: S.L.T.L., X.M.C., G.J.B., Z.V.J., R.C.L.

