## Supplemental Methods for "Natural variation in the oxytocin receptor gene predicts social observation in female prairie voles"

### Random Forest Classifier (RFC) Features

The mathematical definitions of the features used for behavioral analysis are provided below:

Let $c(t) = \left(x_{\text{center}}(t), y_{\text{center}}(t)\right)$ denote the body center and $n(t) = \left(x_{\text{nose}}(t), y_{\text{nose}}(t)\right)$ the nose position at frame $t$, and let $\Delta t$ be a temporal interval.

**Core Features**

Feature 1 – 2, Speed of forward motion of the body center (for each animal):

$$v_{\text{body}}(t) = \frac{\|c(t+\Delta t) - c(t-\Delta t)\|}{2\Delta t} \cdot \cos\left(\theta_{\text{head}}(t) - \theta_{\text{motion}}(t)\right)$$

where $\Delta t = 4$, $\theta_{\text{motion}}(t) = \tan^{-1}\left(\frac{dy}{dx}\right)$, $\theta_{\text{head}}(t) = \tan^{-1}\left(\frac{y_{\text{nose}}(t) - y_{\text{center}}(t)}{x_{\text{nose}}(t) - x_{\text{center}}(t)}\right)$

Feature 3 – 4. Speed of forward motion of the nose (for each animal):

$$v_{\text{nose}}(t) = \frac{\|n(t+\Delta t) - n(t-\Delta t)\|}{2\Delta t} \cdot \cos\left(\theta_{\text{head}}(t) - \theta_{\text{motion}}(t)\right)$$

Feature 5 – 6. Change in head direction (for each animal):

$$\Delta\theta_{\text{head}}(t) = \theta_{\text{head}}(t) - \theta_{\text{head}}(t-1)$$

Feature 7 – 8. Body area with the shoelace formula (for each animal):

$$A(t) = \frac{1}{2}\left|\sum_{k=1}^{6} x_k y_{k+1} - x_{k+1} y_k\right|, \quad x_7 = x_1,\ y_7 = y_1$$

where $(x_k, y_k)$ are the coordinates of six ordered (clockwise or counterclockwise) landmarks on the outline of the body.

Feature 9 – 10. Body aspect ratio (for each animal):

$$R(t) = \frac{\left\|p_{\text{nose}} - p_{\text{anogenital}}\right\|}{\left\|p_{\text{body(left)}} - p_{\text{body(right)}}\right\|}$$

Feature 11 – 12. Relative heading angle to partner's body center (for each animal):

$$\theta_{\text{rel}}(t) = \cos^{-1}\left(\frac{\overrightarrow{v_{\text{head}}} \cdot \overrightarrow{v_{\text{target}}}}{\left\|\overrightarrow{v_{\text{head}}}\right\|\left\|\overrightarrow{v_{\text{target}}}\right\|}\right)$$

where $\overrightarrow{v_{\text{head}}} = n(t) - c(t)$, $\quad \overrightarrow{v_{\text{target}}} = c_{partner}(t) - n(t)$

Feature 13. Inter-animal body center distance:

$$d_{\text{center}}(t) = \|c_A(t) - c_B(t)\|$$

Feature 14. Nose-to-nose distance:

$$d_{\text{nose}}(t) = \|n_A(t) - n_B(t)\|$$

**Normalization**

Features involving speed (Features 1 – 4) or area (Features 7 – 8) were normalized by the individual's median major axis length to account for differences in body size.

$$f_{\text{normalized}}(t) = \frac{f(t)}{\text{median(major axis)}}$$

**Smoothed Features**

We applied Gaussian smoothing to reduce frame-wise jitter using a window of 11 frames centered on $t$:

$$f_{\text{smoothed}}(t) = \sum_{k=-5}^{5} f(t+k) \cdot \frac{1}{\sqrt{2\pi}\sigma} \exp\left(-\frac{k^2}{2\sigma^2}\right)$$

Smoothed versions (Features 15 – 26) were computed for:

Feature 15 – 16 from Feature 1 – 2: $v_{\text{body}}^{\text{smooth}}(t)$

Feature 17 – 18 from Feature 3 – 4: $v_{\text{nose}}^{\text{smooth}}(t)$

Feature 19 – 20 from Feature 7 – 8: $A^{\text{smooth}}(t)$

Feature 21 – 22 from Feature 9 – 10: $R^{\text{smooth}}(t)$

Feature 23 – 24 from Feature 11 – 12: $\theta_{\text{rel}}^{\text{smooth}}(t)$

Feature 25 from Feature 13: $d_{\text{center}}^{\text{smooth}}(t)$

Feature 26 from Feature 14: $d_{\text{nose}}^{\text{smooth}}(t)$

### Linear Mixed Effect Model Analysis of Social Preference Tests (SPT)

To account for the random effect of individual variance, data from the SPT (**Figs 1D, 6A-C**) were additionally analyzed using linear mixed-effects (LME) models with an ANOVA to assess fixed effects. These were implemented in MATLAB (fitlme), with subject included as a random effect. When residual diagnostics indicated deviations from model assumptions such as heteroscedasticity, outcome variables were log-transformed prior to analysis, and model assumptions were re-evaluated to confirm adequacy.

Table S1: Figure 1D LME results

| GLM Summary Table | Estimate ± SE | F (df1, df2) | p-value | partial η² | Sig. |
|---|---|---|---|---|---|
| (Intercept) | 33.023 ± 8.337 | 15.689 (1, 36) | 3.38e-04 | 0.30352 | *** |
| Genotype | -7.804 ± 11.791 | 0.43812 (1, 36) | 0.51225 | 0.012024 | |
| Social zone | 15.459 ± 11.791 | 1.7189 (1, 36) | 0.19813 | 0.045572 | |
| Genotype × Social zone | 6.251 ± 16.675 | 0.14055 (1, 36) | 0.70993 | 0.0038891 | |

Table S2: Figure 6A LME results

| GLM Summary Table | Estimate ± SE | F (df1, df2) | p-value | partial η² | Sig. |
|---|---|---|---|---|---|
| (Intercept) | 7.414 ± 1.290 | 33.048 (1, 36) | 1.51e-06 | 0.47863 | *** |
| Genotype | -3.246 ± 1.824 | 3.1679 (1, 36) | 0.083542 | 0.08088 | |
| Social zone | -2.928 ± 1.824 | 2.5773 (1, 36) | 0.11715 | 0.066808 | |
| Genotype × Social zone | 6.460 ± 2.579 | 6.2719 (1, 36) | 0.016935 | 0.14837 | * |

Table S3: Figure 6B LME (log-transformed) results

| GLM Summary Table | Estimate ± SE | F (df1, df2) | p-value | partial η² | Sig. |
|---|---|---|---|---|---|
| (Intercept) | 0.162 ± 0.080 | 4.1376 (1, 36) | 0.049359 | 0.10309 | * |
| Genotype | 0.163 ± 0.104 | 2.4753 (1, 36) | 0.12439 | 0.064335 | |
| Social target | 0.220 ± 0.113 | 3.8024 (1, 36) | 0.059001 | 0.095531 | |
| Genotype × Social target | -0.398 ± 0.146 | 7.3863 (1, 36) | 0.010043 | 0.17024 | * |

Table S4: Figure 6C LME (log-transformed) results

| GLM Summary Table | Estimate ± SE | F (df1, df2) | p-value | partial η² | Sig. |
|---|---|---|---|---|---|
| (Intercept) | 1.144 ± 0.111 | 106.89 (1, 34) | 4.9383e-12 | 0.10309 | *** |
| Genotype | 0.693 ± 0.148 | 21.773 (1, 34) | 4.6341e-05 | 0.064335 | *** |
| Context | -0.982 ± 0.148 | 43.748 (1, 34) | 1.3879e-07 | 0.095531 | *** |
| Genotype × Context | -0.530 ± 0.204 | 6.7414 (1, 34) | 0.013815 | 0.17024 | * |
